# Context-aware geometric deep learning for protein sequence design

**DOI:** 10.1101/2023.06.19.545381

**Authors:** Lucien F. Krapp, Fernando A. Meireles, Luciano A. Abriata, Matteo Dal Peraro

## Abstract

Protein design and engineering are evolving at an unprecedented pace leveraging the advances of deep learning. Current models nonetheless cannot natively consider non-protein entities within the design process. Here we introduce a deep learning approach based solely on a geometric transformer of atomic coordinates that predicts protein sequences from backbone scaffolds aware of the restraints imposed by diverse molecular environments. This new concept is anticipated to improve the design versatility for engineering proteins with desired functions.

---

Designing proteins *de novo* to engineer their properties for functional tasks is a grand challenge with direct implications for biology, medicine, biotechnology, and materials science. While physics-based approaches have had success in finding amino acid sequences that fold to a given protein structure, deep learning methods have recently brought a dramatic acceleration by enhancing the design success rates and versatility. Among the most recent and notable examples, ProteinMPNN, based on an encoder-decoder neural network, is able to generate protein sequences experimentally proven to fold as intended^1,2^. More recently, coupled with denoising diffusion probabilistic models for the generation of protein backbones, ProteinMPNN in RFdiffusion has shown remarkable success.^3^ In addition, ESM-IF1, based on a protein language model, is capable of generating highly diverse proteins well outside the known universe of natural sequences^4,5^. The model has also recently found experimental validation reporting a very high success rate^6^. Deep learning approaches are however pervasive in the field finding broad application in several protein design tasks^7–10^, like for example MaSIF which specializes in the design of protein interactions via learned protein surface fingerprints^11,12^.

Although these models can natively handle multiple protein chains in their inputs, and as such they can design the sequences of interacting proteins, they cannot natively consider non-protein entities within the design process, which hampers their versatility and limit their spectrum of application. Here, to address this limitation, we introduce CARBonAra (namely, Context-aware Amino acid Recovery from Backbone Atoms and heteroatoms), a new protein sequence generator model based on our recent Protein Structure Transformer (PeSTo^13^), a geometric transformer architecture that operates on atom point clouds. Representing molecules uniquely by element names and coordinates, PeSTo’s transformer can be applied to and predict protein interfaces with virtually any kind of molecules, either other proteins, nucleic acids, lipids, ions, small ligands, or cofactors. Based on the same architecture, trained uniquely on structural data available on the PDB, CARBonAra predicts the amino acid confidence per position from a backbone scaffold alone or complexed by any kind of non-protein molecules. The model uses geometrical transformers to encode the local neighbourhood of the atomic point cloud using the geometry and atomic elements. It encodes the interactions of the nearest neighbours and employs a transformer to decode and update the state of each atom. By pooling the atom states from the atomic to the residue level and decoding them, the model predicts multi-class residue-wise amino acid confidences (**Figure 1a** and **Methods**). CARBonAra thus provides a potential sequence space that can be refined through the incorporation of specific constraints, such as a molecular context critical to the protein’s function, a particular objective, or provided allowed conformations. CARBonAra offers a novel level of flexibility in protein design by recognizing and incorporating any molecular context into its sequence predictions. This distinctive capability of our method expands therefore the scope of applications in the field of protein design.

**Figure 1.**
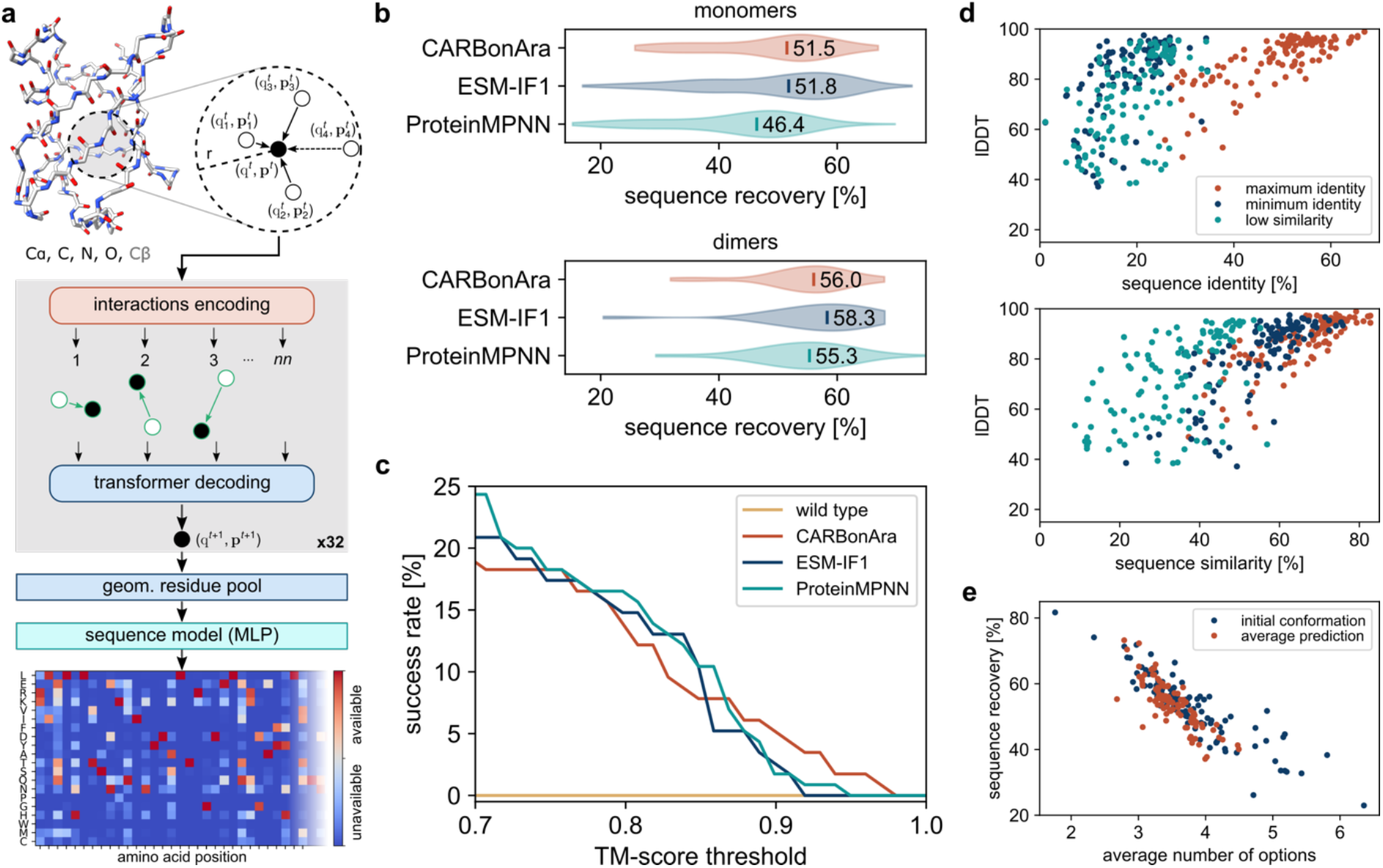
CARBonAra architecture and comparison with other state-of-the-art methods. (**a**) The model applies multiple geometric transformer operations to the coordinates and atom element of a backbone scaffold with added virtual Cβ to predict the amino acid confidence at each position in the sequence. (**b**) Comparison of the sequence recovery of different methods for monomers and dimers with indicated median sequence recovery. (**c**) Percentage of AlphaFold predicted structures, in single sequence mode, above a TM-score threshold. (**d**) Local Distance Difference Test (lDDT) of AlphaFold predicted structures against scaffold monomers from sequences generated using CARBonAra with, as objective, maximum sequence identity, minimum sequence identity, and low sequence similarity. (**e**) Comparison of the sequence recovery between the predicted sequence on crystal structures and the consensus sequence predictions derived from 500 frames sampled from 1 μs molecular dynamics simulations for 80 monomers.

CARBonAra performs on par with state-of-the-art methods like ProteinMPNN and ESM-IF1 for sequence prediction of isolated proteins or protein complexes (**Figure 1b**), while having a similar computational cost taking only a few seconds per run (∽3 seconds). Our method achieves a median sequence recovery rate of 51.3% for protein monomer design and 56.0% for dimer design when reconstructing protein sequences from backbone structures. Moreover, the success rate of the generated sequences using AlphaFold in single-sequence mode is commendable, especially in generating structures with a TM-score above 0.9 (**Figure 1c**). We observed that the model is able to learn the tighter amino acid packing at protein cores thus resulting in higher recovery rates and fewer amino acid possibilities for buried amino acids (**Supplementary Figure 1a-c**). As such, CARBonAra confidently recovers core amino acids while demonstrating greater flexibility on the protein’s surface, unless additional functional or structural constraints are provided.

In contrast to other methods, CARBonAra uses multi-class amino acid predictions that generate a space of potential sequences, opening various possibilities for sequence sampling. For example, one can tailor sequences to meet specific objectives, such as achieving maximal or minimal sequence identity, or low sequence similarity in order to design unique sequences with a specific fold (**Figure 1d** and **Supplementary Figure 2**, see also Methods). An informative way to refine the sequence space uses dynamics as a constraint. By applying CARBonAra to structural trajectories from molecular dynamics (MD) simulations, we were able to improve sequence recovery, especially in cases that previously showed low recovery rates (**Figure 1e**). Simultaneously, we observed a reduction in the number of possible amino acids predicted per position. This further limit the sequence space and could enable the design of targeted structural conformations.

More importantly, leveraging PeSTo’s architecture, this model has the new ability to perform protein sequence prediction conditioned by a specific non-protein molecular context. On a test set similar to the one used for PeSTo, we show that the overall structure median sequence recovery increased from 54% to 58% (**Supplementary Figure 3**) when an additional molecular context is provided. In particular, CARBonAra achieves median sequence recovery rates at the interface of 56% when protein interacting partners are considered and 55% when nucleic acids are used as interfacial restraints, providing a significant improvement over predictions without context (**Figure 2a**). Similarly, the recovery rate at the interface improved significantly if small-molecule entities such as ions (67%), lipids (57%), and ligands (60%) are included (**Figure 2a**). Including these molecules boosts sequence recovery in their surroundings, and reduces the number of amino acid possibilities to sample from (**Figure 2b**). This shows CARBonAra’s power to properly craft the residue types required for the binding of specific molecules.

**Figure 2.**
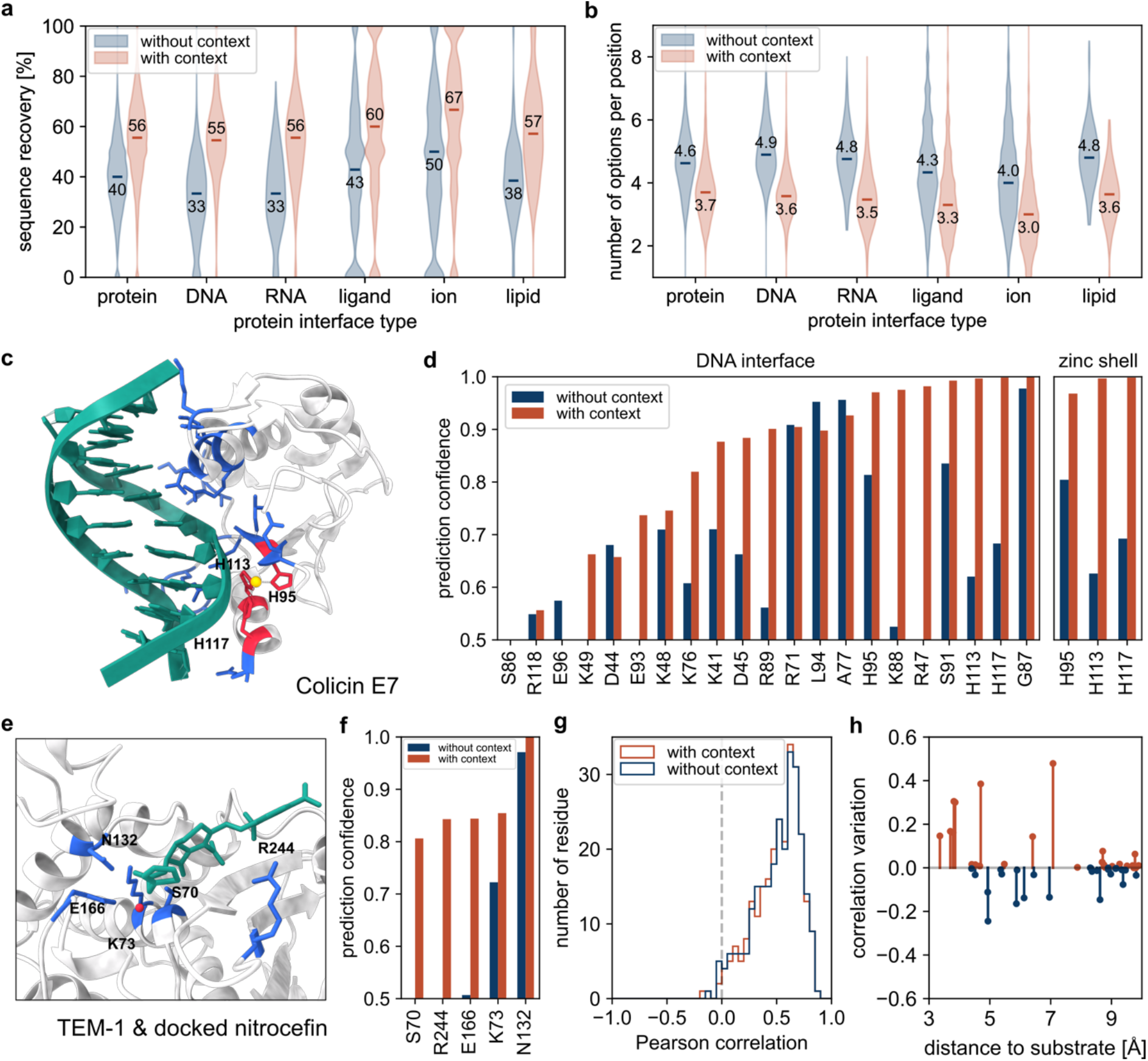
Context-aware amino acid recovery extends to various biomolecules. (**a**) Sequence recovery at the interface (residues within 5 Å) without and with proteins, nucleic acids, ligands, ions, and lipids binders. (**b**) Number of predicted possible amino acids per position at the interface (residues within 5 Å) without and with proteins, nucleic acids, ligands, ions, and lipids binders (considering a confidence prediction threshold of 0.5). (**c**) Colicin E7 endonuclease domain in complex with DNA and a zinc ion (PDB: 1ZNS). The protein-DNA interface (residues within 4 Å) is highlighted in blue. The protein-zinc shell is highlighted in red (residues within 3 Å). (**d**) Estimated accurate prediction probability for the scaffold amino acids at the protein-DNA interface and the protein-zinc shell with and without the presence of DNA and zinc. (**e**) Nitrocefin docked in the active site of the β-lactamase TEM-1 (PDB: 1BT5). Relevant residues for substrate recognition and hydrolysis are shown in blue, nitrocefin in green, and the catalytic water molecule in red. (**f**) Prediction confidence with and without the substrate for the relevant amino acids for binding. (**g**) Correlation of the predictions with deep sequencing analysis of TEM-1. (**h**) Correlation variation by adding the context (nitrocefin and catalytic water) for the amino acids close (in Cβ distance) to the substrate.

An exemplary case to illustrate the power of this approach is the endonuclease domain of ColE7, which interacts with duplex DNA in a zinc-dependent manner^14^. The sequence recovery rate obtained by CARBonAra showed a significant increase from 29% to 52% at the metal and DNA interfaces when the zinc ion or the 12-bp DNA duplex was included as resolved in the native structure (**Figure 2d**). Thus, imposing the presence of non-protein interacting interfaces can enhance the sequence recovery rate significantly, also with respect to predictions done by ProteinMPNN (24%) and ESM-IF1 (43%) (**Supplementary Table 1**). Interestingly, when a non-native molecular context is provided such as a larger ion (e.g., calcium) the sequence recovery rate decreased (**Supplementary Figure 4**). Thus, the predicted amino acid confidence of an ion pocket is widely dependent on the given context, as illustrated also for the metallo β-lactamase BJP-1 (**Supplementary Figure 5**).

Relevant for enzyme design is the possibility to design sequences under the restraints provided by a desired substrate or high-affinity ligand. To test this case, we explored CARBonAra’s ability to predict the sequence of a TEM-1 β-lactamase-like enzyme when the native context at the active site is provided (**Figure 2e**). Without context, the catalytic S70 and substrate binding R244 are never predicted positively (confidence of 0.39 and 0.11 respectively, **Figure 2f**), however, when the prediction is done with a β-lactam (here nitrocefin) docked at the catalytic pocket, the catalytic triad S70, K73, and E166, along with key residues necessary to β-lactam binding (i.e., N132, R244) all have a high prediction confidence (> 0.8) and low ranking (top 2) (**Supplementary Figure 6**). Importantly, in this case, the sequence recovery is maximal when also the catalytic water is considered, hinting at a very high sensitivity for the molecular context.

Given that TEM-1 β-lactamase has been widely studied, we took the occasion to probe what information CARBonAra’s residue-wise amino acid probabilities provide when compared to experimental data. We correlated the estimated probabilities to the residue-wise amino acid probabilities measured experimentally through deep sequencing of a saturated mutagenesis library of the TEM-1 β-lactamase^15^ (**Figure 2g**). We observed an average correlation of 0.51 ± 0.21 for CARBonARa with deep sequencing data, which is similar to the correlation between the deep sequencing data with the multiple sequence alignment of this enzyme’s family (0.52 ± 0.22). This shows that CARBonARa’s estimated probabilities can capture functional sequence variability, a central topic in the realm of protein evolution^16,17^. Moreover, we observed that adding the context to the active site of TEM-1 (i.e. docked nitrocefin and the catalytic water) improved the correlation locally (i.e. for amino acids within 5 Å) but also affects the predictions of amino acids further away (up to 10 Å). These results hint at the possibility to use CARBonAra for the study of the effect of a specific context locally as well as their long-range influences (**Figure 2h**).

In summary, CARBonAra is a novel, structure-centric way to predict protein sequences given a backbone geometry. Leveraging a geometric transformer architecture that can handle any molecular context, CARBonAra can optimize the sequence prediction by considering not only interacting protein partners but also nucleic acids, lipids, small molecules, and ions. This new concept, coupled with the possibility to sample backbone conformations using modern diffusion models, opens exciting new possibilities for the functional design of novel protein-based materials and therapeutics.

## Availability

All datasets, methods and the CARBonAra source code presented in this work will be released upon publication at https://github.com/LBM-EPFL/CARBonARa.

## Acknowledgments

The Swiss National Science Foundation is acknowledged for supporting this work (grant number 205321_192371 to MDP). We also acknowledge the Swiss National Supercomputing Centre (CSCS) for the generous computing time allocation used to run molecular dynamics simulations.

## Author contributions

LFK and MDP conceived and designed the research project. LFK designed and implemented the CARBonAra code. LFK, FAM, LAA, and MDP analysed the data. LFK, FAM, LAA, and MDP wrote the paper.

## Supplementary Information

**Supplementary Algorithm 1.**
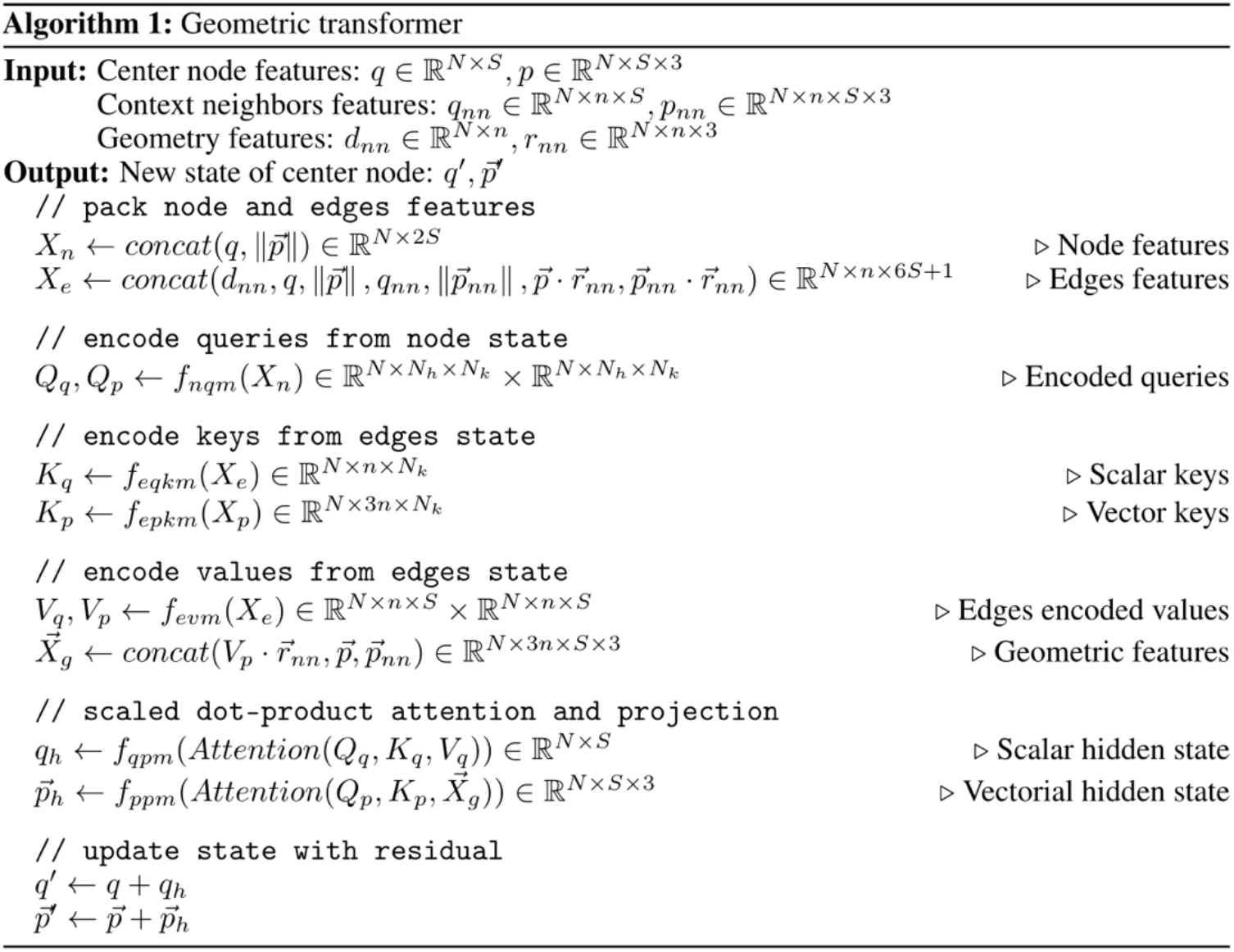
CARBonAra geometric transformer. Based on the PeSTo architecture, each geometric transformer is composed of 5 neural networks of 3 layers with an exponential linear unit (ELU) activation function. The characteristic dimensions are the number of atoms (N), the state size (S), the number of nearest neighbors (nn), the dimension of the embedding for the keys (Nk) and the number of attention heads (Nh). The neural networks haven a flat architecture with hidden layers width equal to the input and output state size (S). The multi-layers perceptrons (MLP) are the node query model (fnqm), encoding scalar key model (feqkm), encoding vector key model (fepkm), encoding value model (fevm), and scalar state projection model (fqpm). The vectorial hidden state is projected over the attention heads with a weighted sum (Wppm) to preserve the rotation equivariance of the operation. The output vector state belongs to the span of the geometry and vector states.

**Supplementary Table 1.**
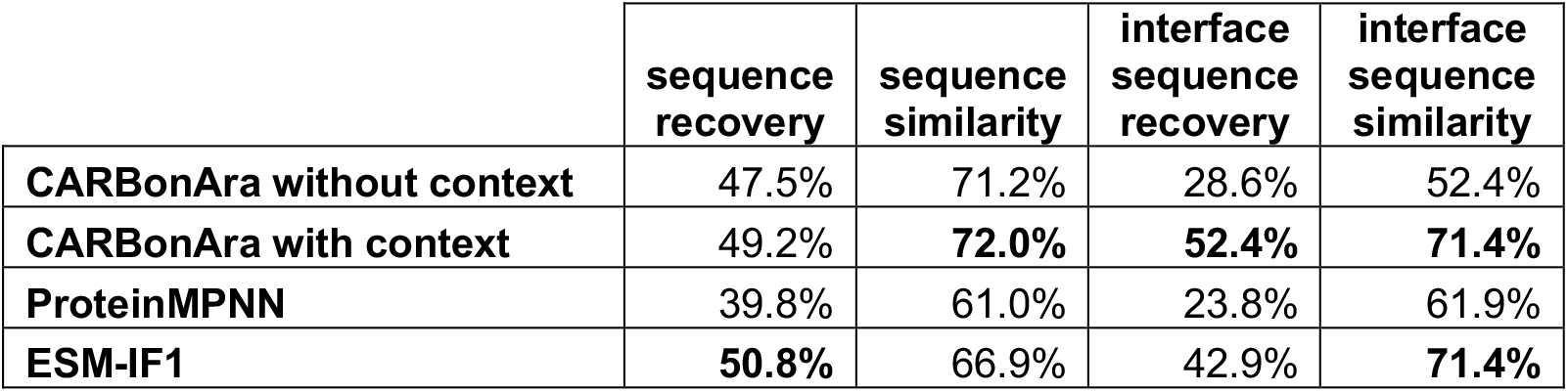
Method comparison with and without DNA bound on Colicin E7. Sequence recovery and similarity for the whole structure and at the interface (residues within 4 Å) with and without DNA of Colicin E7.

|  | sequence recovery | sequence similarity | interface sequence recovery | interface sequence similarity |
| --- | --- | --- | --- | --- |
| **CARBonAra without context** | 47.5% | 71.2% | 28.6% | 52.4% |
| **CARBonAra with context** | 49.2% | **72.0%** | **52.4%** | **71.4%** |
| **ProteinMPNN** | 39.8% | 61.0% | 23.8% | 61.9% |
| **ESM-IF1** | **50.8%** | 66.9% | 42.9% | **71.4%** |

**Supplementary Figure 1.**
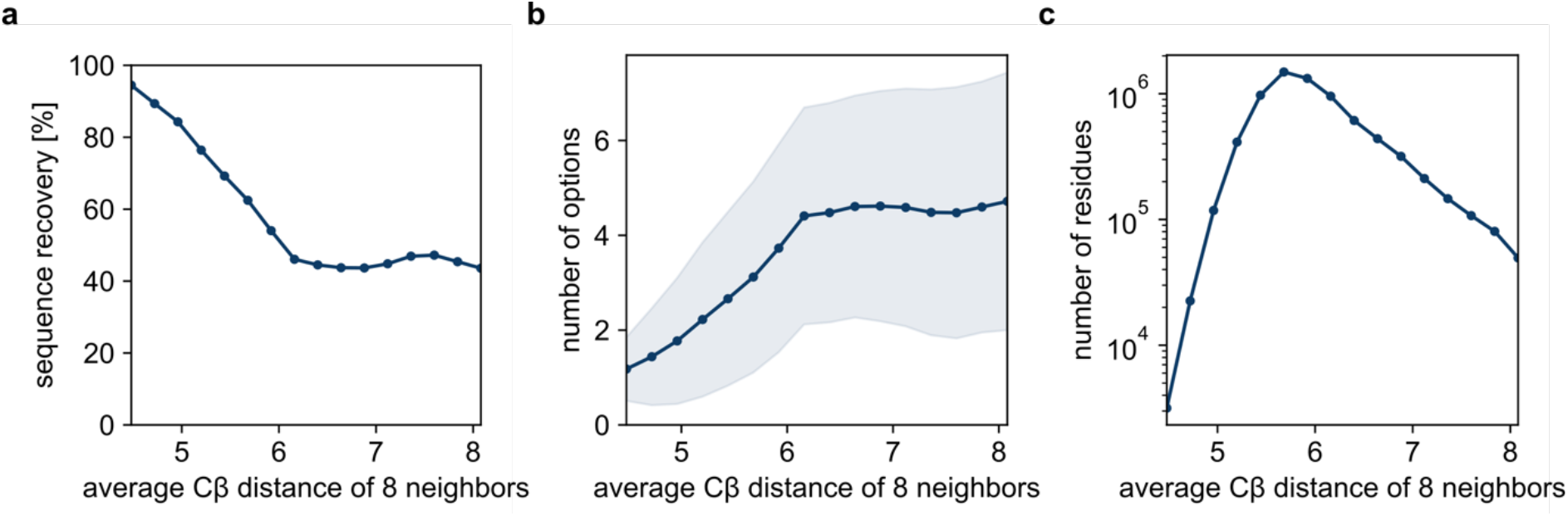
Analysis of buried against surface amino acids. (**a**) Sequence recovery, (**b**) number of predicted options per position and (**c**) number of residues as a function of the average Cβ distance of the 8 nearest neighbours (18866 structures from the testing dataset).

**Supplementary Figure 2.**
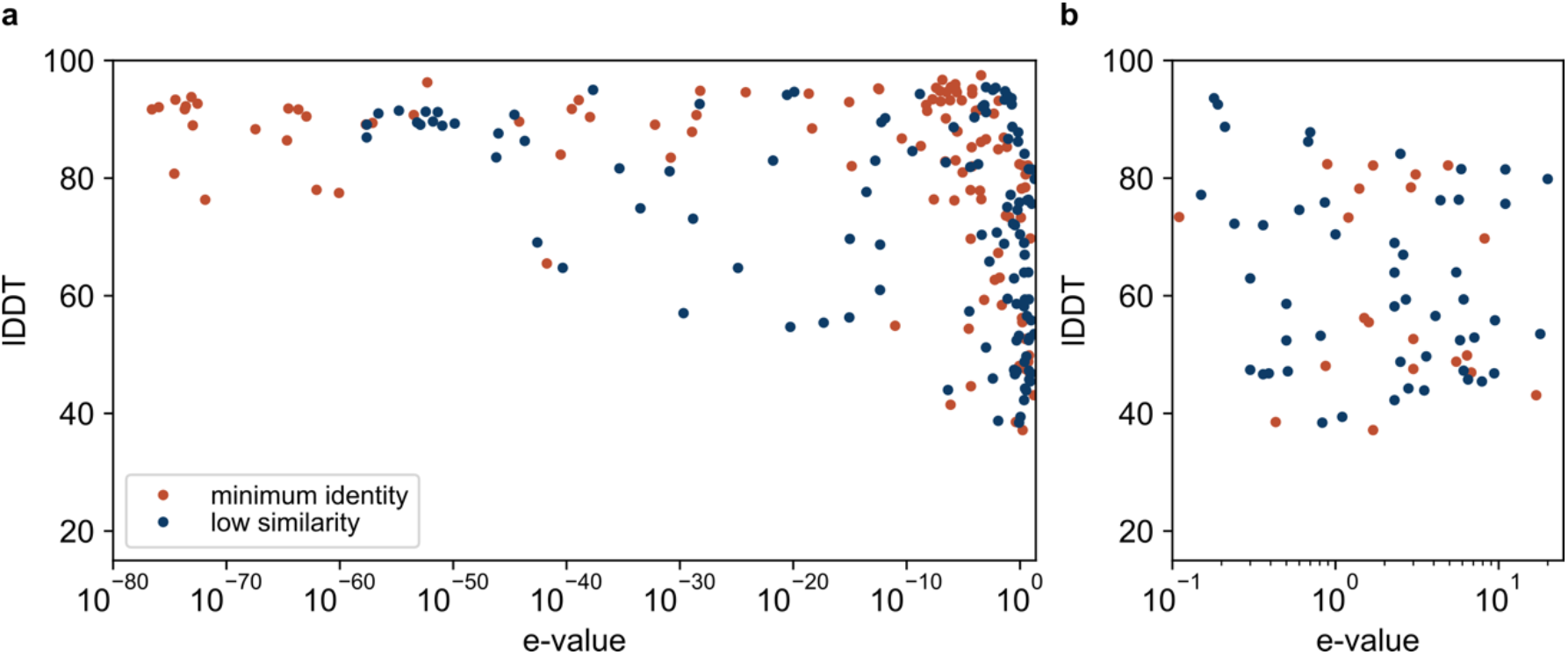
Analysis of the generated sequences. (**a**) lDDT of the AlphaFold predicted structures as a function of the expect value (E-value) of the generated sequences. (**b**) Close up on the generated sequences with a high E-value.

**Supplementary Figure 3.**
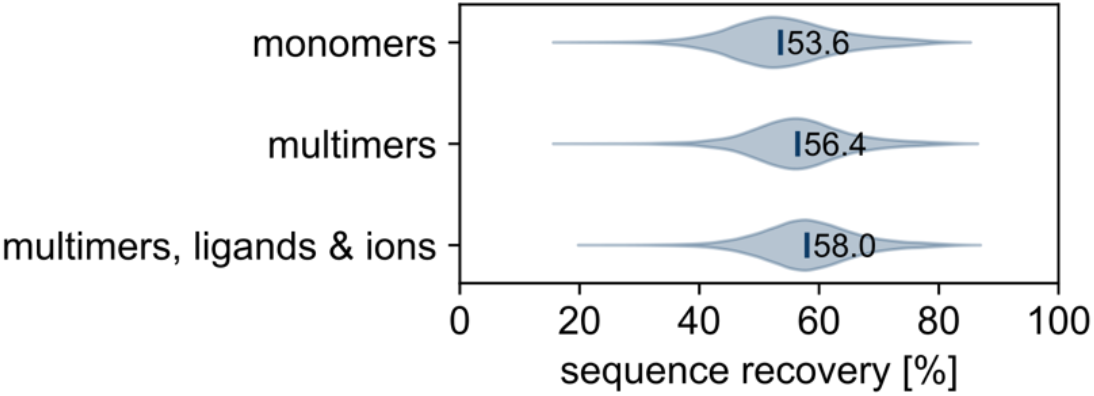
Benchmark of different use cases. Sequence recovery distribution for systems of monomers, multimers and any biomolecules (18866 structures from the testing dataset). The median sequence recovery is indicated for each case.

**Supplementary Figure 4.**
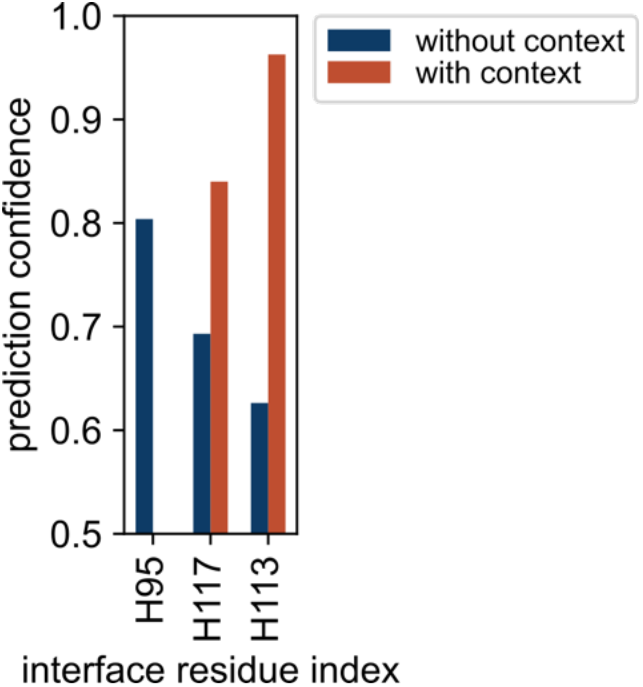
Effect of changing the ion type on the prediction. The prediction confidence for the three most important amino acids for ion binding in the case where the zinc ion of Colicin E7 is replaced with a calcium ion.

**Supplementary Figure 5.**
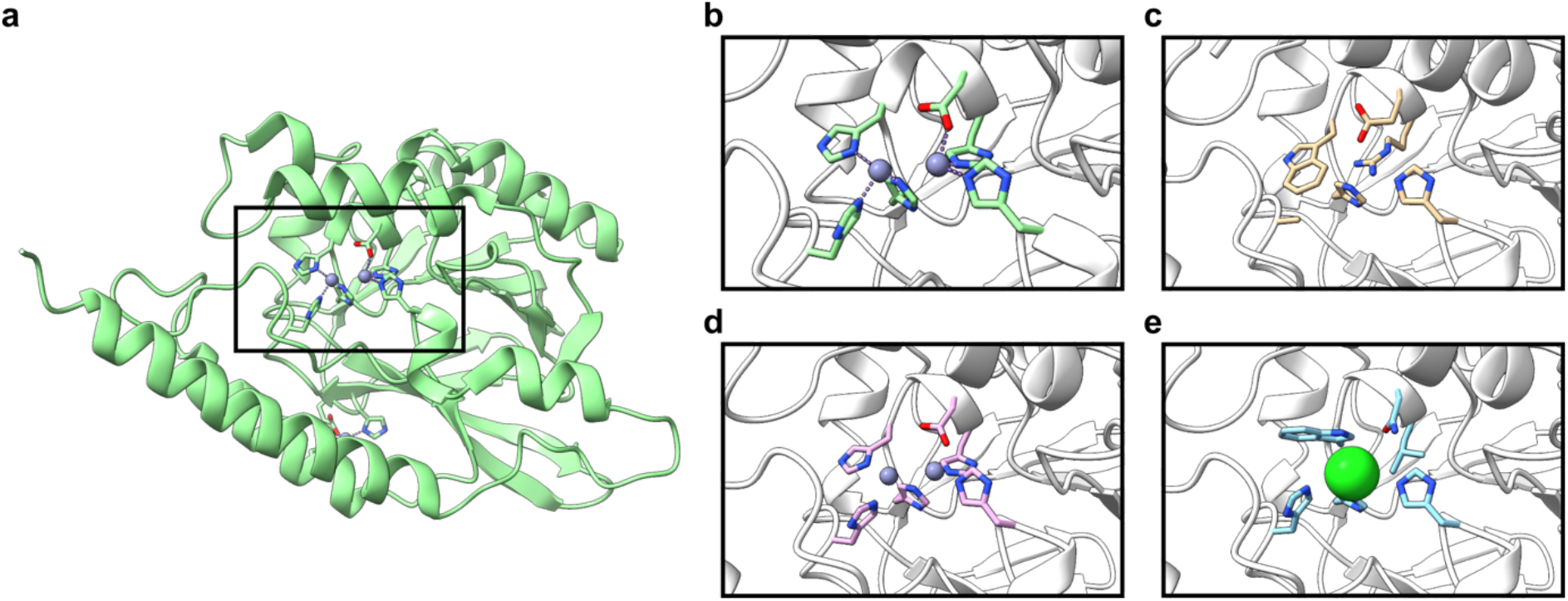
Effect of the ion context on the optimal predicted sequence in the case of a metallo β-lactamase zinc binding pocket. (**a**) Metallo β-lactamase structure with a pocket containing two zinc ions (PDB ID: 3LVZ). (**b**) WT pocket of the metallo β-lactamase. Pocket of an AlphaFold predicted structure with a designed sequence applied to the scaffold structure without zinc ions (**c**), containing the original zinc ions (**d**) and containing a manually placed chloride ion (**e**).

**Supplementary Figure 6.**
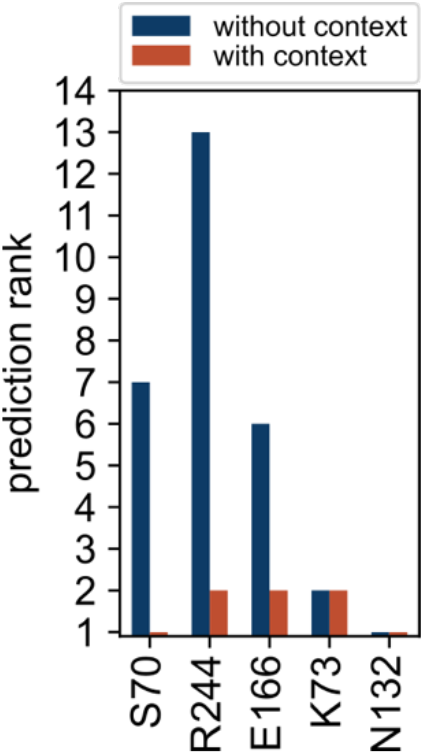
Effect of the docked nitrocefin and catalytic water in TEM-1 on the prediction ranking. Rank of the prediction from maximum to minimum confidence for the 5 important amino acids at the pocket without and with the docked nitrocefin and catalytic water.

**Supplementary Figure 7.**
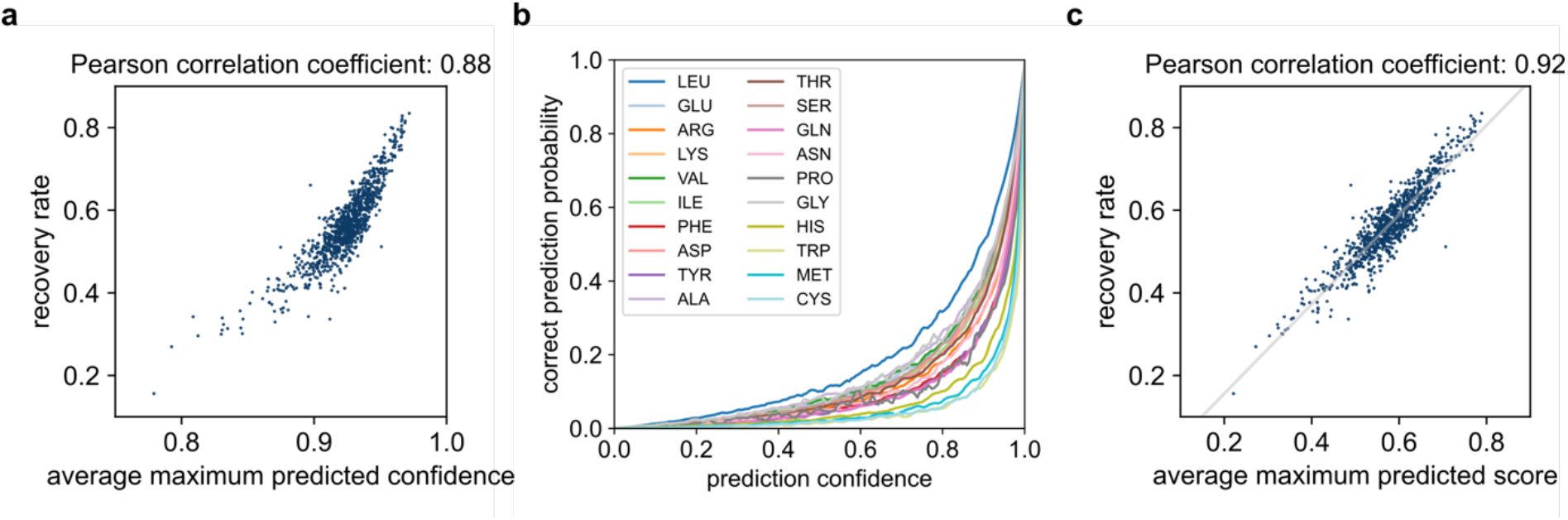
Prediction confidence analysis. (**a**) Recovery rate as a function of the average maximum prediction score (943 structures from the testing dataset). (**b**) Relationship between prediction confidence and the prediction accuracy for each amino acid type (4096 subunits from the training dataset). (**c**) Rescaling prediction score into a prediction confidence correlated with the probability to be correct (943 structures from the testing dataset).

